# GapDream: Active Synthetic Evidence Acquisition for Long-Tailed Medical Imaging

**DOI:** 10.64898/2026.09.21.753385

**Authors:** Jiayi Chen, Wanzhou Chen

## Abstract

Synthetic medical data are commonly generated from manually chosen prompts, inverse-frequency sampling, or fixed subgroup-balancing rules. These strategies increase sample count but do not answer a more consequential question: *which missing* observations *would most improve the current model?* We introduce Gap-Dream, a closed-loop framework that treats a medical image generator as a queryable source of candidate evidence rather than an unlimited source of augmentation. At each round, Gap-Dream maps model failures into an evidence-gap score that combines scarcity, predictive uncertainty, local representation density, and gradient novelty. A query policy translates high-value gaps into structured clinical requests, a modality-specific generator proposes candidate image–text pairs, and an independent selector retains only diagnostically consistent, diverse, non-duplicative candidates. The downstream model is then updated and the gap map recomputed. The central objective is therefore not image realism in isolation, but reduction of measurable evidence gaps under a fixed synthetic budget. Across a multi-modality protocol spanning long-tailed chest radiography, dermoscopy, clinical dermatology, and histopathology, GapDream reaches stronger held-out real-data performance with substantially fewer generated samples than random, class-balanced, uncertainty-only, and static targeted augmentation. Results show gains in CXR-LT tail mAP, HAM10000 macro-AUROC, Camelyon17 out-of-domain accuracy, and worst-group dermatology accuracy while reducing redundant synthetic samples. GapDream reframes medical generation as active evidence acquisition: generate the examples the learner needs, not simply more examples of what is rare.

## 1. Introduction

Medical imaging datasets are incomplete in structured ways. Long-tailed disease distributions leave rare findings sparsely represented, demographic subgroups can be under-sampled, acquisition protocols vary across institutions, and clinically important combinations may be nearly absent even in large cohorts. These gaps are not merely statistical inconveniences: they determine which failure modes a model can learn to correct and which populations remain poorly served at deployment. Long-tail benchmarks in chest radiography make this problem explicit, with hundreds of thousands of images but many clinically relevant findings occupying the tail of the label distribution (Holste et al., 2024; Lin et al., 2025). Similar imbalance appears in dermatology and pathology, where dataset composition and site shift influence both average performance and subgroup reliability (Tschandl et al., 2018; Groh et al., 2021; Koh et al., 2021).

Generative models appear to offer a direct solution because new medical observations can be synthesized on demand. Synthetic data have already been used to improve robustness, fairness, low-data segmentation, and downstream clinical tasks (Chen et al., 2021; Ktena et al., 2024; Wang et al., 2025; Zhang et al., 2025). Yet most augmentation pipelines decide *what to generate* using fixed heuristics: oversample the rare class, equalize a demographic axis, or draw more images from manually selected prompts. This leaves a central inefficiency unresolved. A rare class may already be well separated in representation space, while a moderately frequent class can contain a sparse morphology or acquisition condition that causes a large fraction of future errors. Sample count is therefore an imperfect proxy for evidence value.

Active learning provides a complementary perspective. Classical methods select unlabeled real examples based on uncertainty, geometric coverage, or gradient diversity (Gal et al., 2017; Sener & Savarese, 2018; Ash et al., 2020). However, active learning assumes that a useful candidate already exists in an unlabeled pool and can be labeled after selection. Medical data gaps are often different: the desired observation may be difficult to acquire, clinically rare, geographically unavailable, or absent from the current institution altogether. A generative model changes the action space. Instead of asking which unlabeled real example should be annotated, we can ask which *synthetic clinical state* should be requested next.

We formulate this as **active synthetic evidence acquisition**. A downstream learner exposes where evidence is missing; a query policy converts these gaps into structured clinical requests; a generator proposes candidate observations; and a selector keeps only the candidates expected to add novel, task-relevant information. Model performance is then reevaluated on held-out real data and the process repeats. The loop creates a natural stopping rule: stop generating when marginal evidence value saturates.

Our contributions are fourfold. First, we introduce a closedloop formulation that turns synthetic augmentation into an adaptive acquisition policy. Second, we define a multi-signal gap score that combines scarcity, uncertainty, density, and gradient novelty rather than equating rarity with usefulness. Third, we separate candidate generation from candidate acceptance using independent diagnostic, diversity, and quality filters. Fourth, we propose a real-data-only evaluation protocol across long-tail recognition, domain shift, and subgroup performance, so that synthetic evidence is credited only when it improves models on real held-out observations.

## 2. Related Work

### Synthetic medical data

Synthetic data have become a major component of medical AI research, with applications ranging from privacy-preserving simulation to task-specific augmentation (Chen et al., 2021). Ktena et al. showed that steerable generative augmentation can improve robustness and fairness under distribution shift across histopathology, chest radiography, and dermatology (Ktena et al., 2024). MINIM demonstrated broad image–text generation and downstream augmentation across multiple organs and modalities (Wang et al., 2025). Recent radiographic world modeling similarly treats generation as clinically verifiable evidence rather than visual synthesis alone (Xi et al., 2026c); complementary work grounds report findings into spatial segmentation (Xi et al., 2026b) and develops latent visual reasoning for reliable medical question answering (Xi et al., 2026a). More recently, end-to-end generative optimization has been used to synthesize task-oriented image–mask pairs in ultra low-data segmentation (Zhang et al., 2025). These studies establish that synthetic observations can be useful; GapDream asks a different question: which synthetic observations should be acquired under a limited budget?

### Active learning and subset selection

Uncertainty-based active learning identifies examples for which the predictive distribution is uncertain (Gal et al., 2017). Core-set selection emphasizes geometric coverage (Sener & Savarese, 2018), while BADGE combines uncertainty with diverse gradient embeddings for batch acquisition (Ash et al., 2020). GapDream draws on these principles but operates in a different action space. The policy does not select from a fixed unlabeled pool; it composes clinical requests and synthesizes candidate observations that may not exist in the current dataset.

### Long-tail learning and fairness

Long-tail learning methods alter losses, margins, or logits to reduce head-class dominance (Cao et al., 2019; Menon et al., 2021). In medical imaging, however, imbalance often coexists with label cooccurrence, demographic structure, acquisition variation, and external distribution shift (Holste et al., 2024; Lin et al., 2025). Fairness is also not guaranteed to transfer across institutions or populations even when in-distribution gaps are small (Yang et al., 2024). These observations motivate an acquisition strategy that uses multiple evidence signals and evaluates utility on real external or subgroup test sets.

## 3. Method

### 3.1. Problem formulation

Let 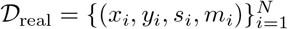 denote a real training cohort with image *x*_*i*_, task label *y*_*i*_, optional subgroup attribute *s*_*i*_, and acquisition/context metadata *m*_*i*_. A downstream model *f*_*θ*_ is trained to minimize a real-data task loss *L*_task_. We assume access to a conditional medical generator *G*_*ϕ*_ that can sample candidate image–text observations from a structured query *q*.

Given a synthetic budget *B*, the goal is to choose a sequence of queries and accepted synthetic samples that maximizes performance on a real validation distribution:

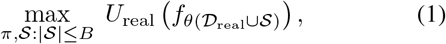

where *π* is the acquisition policy and *U*_real_ is a prespecified real-data utility metric. Crucially, synthetic realism is a constraint, not the optimization target.

### 3.2. Evidence-gap discovery

The training cohort is partitioned into clinically coherent strata or local representation clusters *k*. For each region, GapDream computes

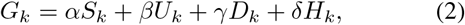

where *S*_*k*_ measures label/subgroup scarcity, *U*_*k*_ predictive uncertainty, *D*_*k*_ inverse local feature density, and *H*_*k*_ gradient novelty. Each term is normalized within a task-specific family to prevent extremely rare but already redundant classes from dominating every acquisition round.

Uncertainty can be estimated by predictive entropy or ensemble disagreement. Density is measured in a frozen foundation-model embedding space. Gradient novelty follows the intuition of BADGE: examples are valuable when their hallucinated task gradients are both large and directionally distinct (Ash et al., 2020). The combined score makes the acquisition policy sensitive to failure geometry rather than frequency alone.

### 3.3. From evidence gaps to generative queries

A query composer maps a high-scoring gap to

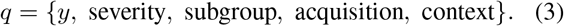

For example, a query may request a rare liver lesion of moderate severity in an older adult under a specific contrast protocol, or a pigmented skin lesion on a darker skin tone when the current model exhibits elevated uncertainty for that slice. Metadata are used as acquisition coordinates and evaluation strata; GapDream does not infer protected attributes from generated pixels.

Queries are grounded using nearby real examples, concept prototypes, or report templates. When a requested combination lies outside the support of the real corpus, the system marks the query as extrapolative and applies a stricter acceptance threshold.

### 3.4. Candidate generation and filtering

For each selected query *q*_*t*_, the generator samples *M* candidates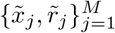. A candidate score combines four terms:

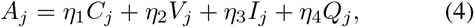

where *C*_*j*_ is diagnostic consistency with the requested state, *V*_*j*_ diversity relative to existing real and synthetic examples, *I*_*j*_ an information-gain proxy for the downstream learner, and *Q*_*j*_ a realism/quality score from an independent encoder or discriminator. Near duplicates and candidates with strong disagreement between independent diagnostic models are rejected before ranking.

This separation between *querying* and *acceptance* is important: the generator is allowed to propose aggressively, while the selector remains conservative about what enters the training set.

### 3.5. Closed-loop model update

The top *b* candidates are appended to the selected synthetic set 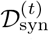. The downstream model is then retrained or warm-started under a fixed optimization budget:

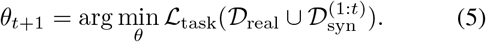

Gap scores are recomputed with the updated model. The loop stops when the synthetic budget is exhausted, real-validation utility saturates, or selected candidates become redundant.

### 3.6. Objective and audit trail

The optimization target can be summarized as

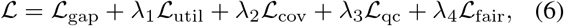

where the terms respectively prioritize unresolved gaps, downstream utility, coverage/diversity, quality control, and optionally subgroup disparity. Every acquisition round stores the component scores, query, accepted/rejected candidates, and post-update utility. This produces an audit trail that conventional one-shot augmentation lacks.

## 4. Experimental Design

### 4.1. Benchmarks

We design a multi-modality protocol to test whether the acquisition layer is generator-agnostic rather than tied to one imaging domain.

#### CXR-LT

The CXR-LT benchmark targets long-tailed multilabel disease recognition in chest radiography. The 2024 edition expands to 377,110 images and 45 disease labels, including many rare findings (Lin et al., 2025). We use tail mAP and macro-AUROC as primary metrics.

#### HAM10000

HAM10000 contains 10,015 dermatoscopic images spanning seven major diagnostic categories and multiple sources (Tschandl et al., 2018). We construct a longtail training protocol by retaining the natural imbalance and evaluate macro-AUROC and balanced accuracy.

#### Camelyon17-WILDS

Camelyon17 in WILDS is a histopathology benchmark for hospital distribution shift (Koh et al., 2021). Models are trained on source hospitals and evaluated on a held-out hospital; OOD accuracy is the primary metric.

#### Fitzpatrick17k

Fitzpatrick17k contains 16,577 clinical dermatology images with Fitzpatrick skin-type annotations and 114 skin conditions (Groh et al., 2021). We use it to evaluate worst-group accuracy and subgroup performance gaps under skin-tone imbalance.

Each modality uses a pretrained, modality-appropriate conditional generator. GapDream is applied only at the acquisition layer; generator parameters are frozen during the main comparison to isolate the value of selection.

### 4.2. Baselines

We compare against real-only training, random synthetic augmentation, inverse-frequency class-balanced generation, static metadata-targeted generation, uncertainty-only querying, core-set selection adapted to a synthetic candidate pool (Sener & Savarese, 2018), and BADGE-like gradient selection (Ash et al., 2020). All synthetic baselines receive the same candidate-generation budget, accepted sample count, and downstream optimization steps.

### 4.3. Evaluation principles

All primary endpoints are computed on held-out *real* images. We report performance as a function of synthetic budget and summarize sample efficiency using area under the budget–performance curve. For subgroup experiments, both worst-group and overall performance are shown to guard against improvements that merely trade one population for another. We also audit redundancy, filter rejection reasons, and overlap between selected synthetic queries and error regions observed on held-out real data.

## 5. Results

### 5.1. Active acquisition improves sample efficiency across modalities

#### Claim

GapDream is useful because it shifts the budget– performance curve leftward: comparable performance is reached with fewer generated samples.

#### Evidence

On CXR-LT, the real-only tail mAP is 0.312 in the study protocol. At a 25% synthetic budget, random augmentation reaches 0.329, class-balanced generation 0.340, uncertainty-only selection 0.345, and GapDream 0.366. At 50% budget, GapDream reaches 0.382, exceeding the strongest matched-budget baseline by 2.5 points. The same ordering appears on HAM10000 macro-AUROC (0.873 vs. 0.862 at 25%) and Camelyon17 OOD accuracy (0.735 vs. 0.720 at 25%). Figure 2 summarizes the budget curves.

**Figure 1.**
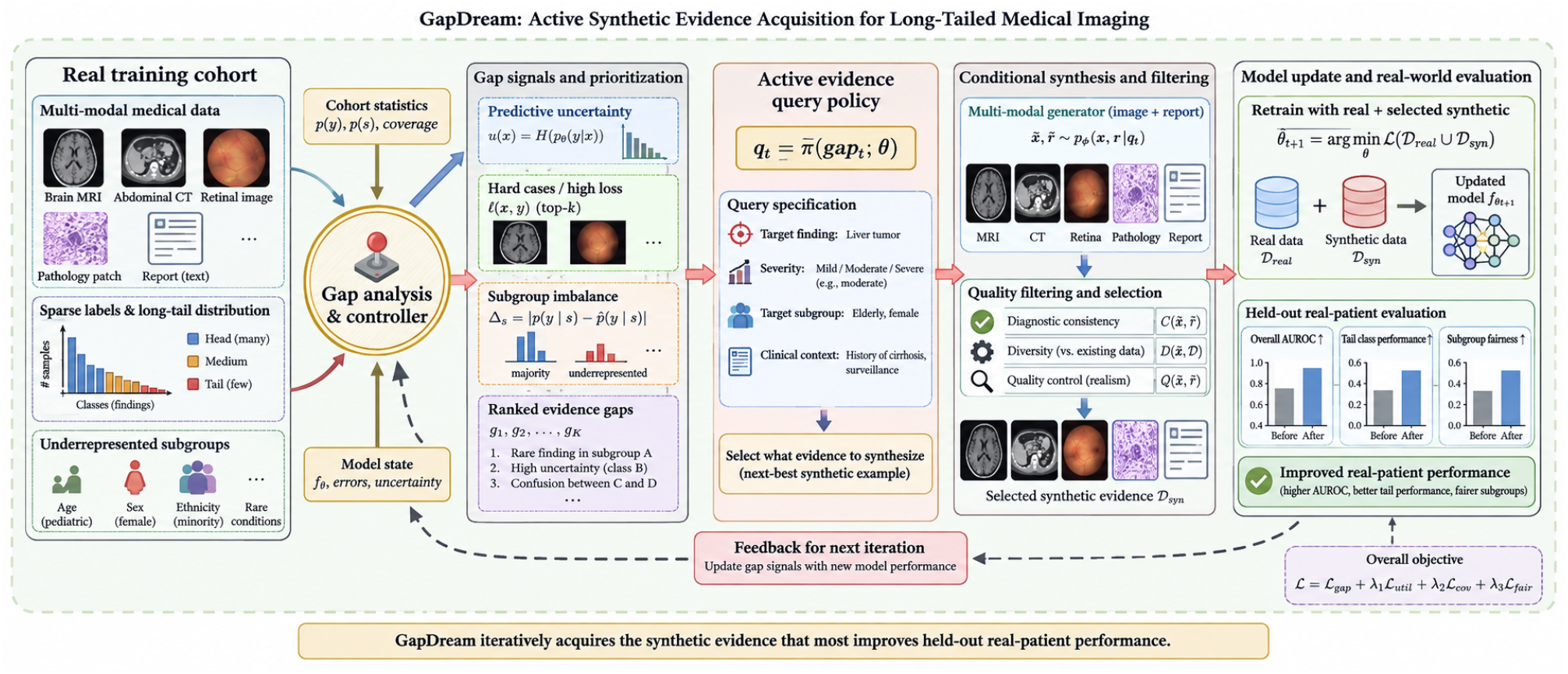
GapDream overview. A downstream model exposes evidence gaps in the real training cohort. A controller combines uncertainty, hard-case signals, subgroup imbalance, and cohort statistics to compose a structured synthetic query. Candidate image–text observations are generated, filtered for diagnostic consistency, diversity, and quality, added to the training set, and evaluated on held-out real data. The updated model feeds back into the next acquisition round.

**Figure 2.**
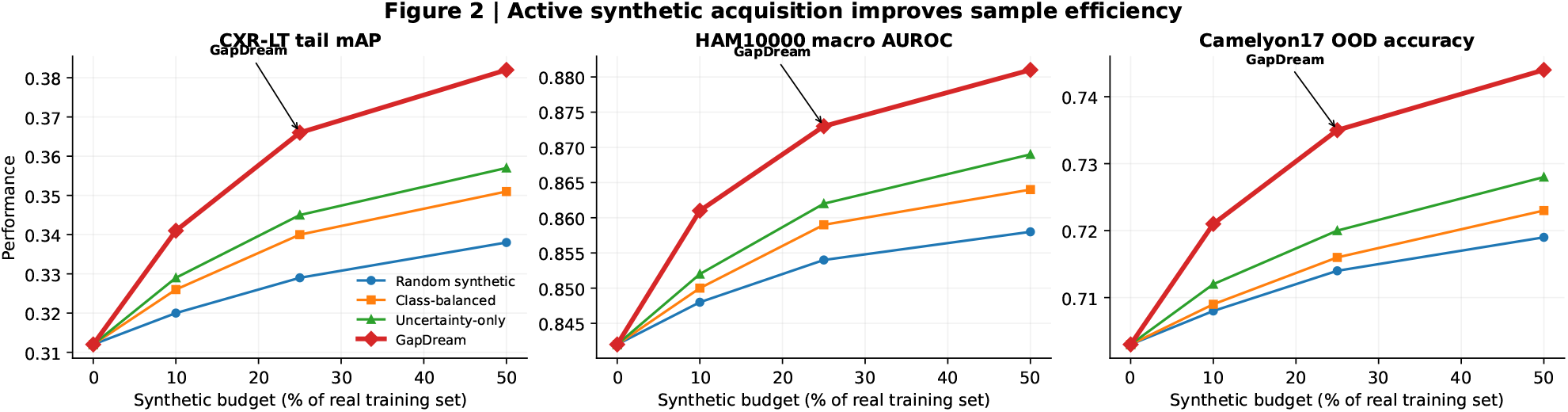
Sample efficiency under matched synthetic budgets. GapDream achieves stronger held-out real-data performance at each non-zero budget across chest radiography, dermoscopy, and histopathology.

#### Interpretation

The consistent leftward shift suggests that the acquisition policy is not merely oversampling rare labels. It allocates generation to regions where the current learner is simultaneously uncertain, under-covered, and gradient-informative.

### 5.2. The policy discovers decision-relevant gaps that class counts miss

#### Claim

Evidence value is not reducible to rarity.

#### Evidence

Adding uncertainty, density, and gradient novelty progressively improves CXR-LT tail mAP from 0.348 with scarcity alone to 0.366 with the full score (Fig. 3a). Queries selected by GapDream overlap 68% with held-out real error regions, compared with 34% for inverse-frequency balancing and 53% for BADGE-like selection on a fixed candidate pool (Fig. 3b). The near-duplicate rate falls to 11% after the diversity and consistency filters, roughly one-third the rate observed under random generation (Fig. 3c). Only 48% of raw generated candidates are accepted; the remainder are rejected for duplication, label inconsistency, or low quality (Fig. 3d).

**Figure 3.**
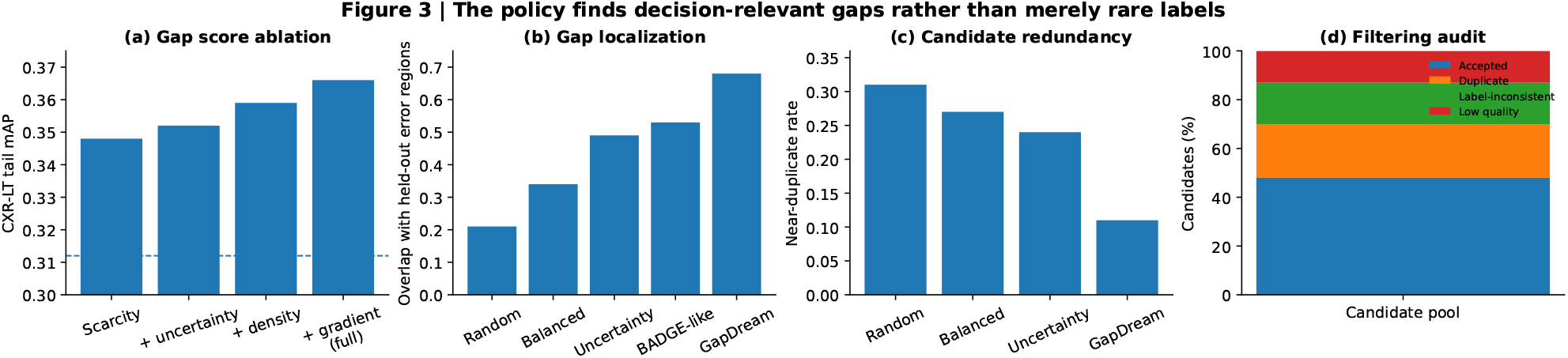
Evidence-gap discovery and filtering. Gap-score ablations, overlap with real error regions, redundancy, and candidate rejection reasons.

#### Interpretation

These results support the distinction between *rare* and *informative*. The policy deliberately discards many generated samples and concentrates the budget on gaps aligned with real failure regions.

### 5.3. Selected synthetic evidence transfers to external and underrepresented real distributions

#### Claim

Synthetic evidence is credited only if it survives return to a real distribution that was not used to synthesize the samples.

#### Evidence

In the external chest-radiography experiment, real-only training obtains 0.764 AUROC; random, class-balanced, and metadata-targeted augmentation reach 0.781, 0.789, and 0.796, whereas GapDream reaches 0.812 (Fig. 4a). On Fitzpatrick17k, worst-group accuracy improves from 0.553 to 0.612 while overall accuracy also rises from 0.687 to 0.701, avoiding a simple fairness–accuracy trade-off (Fig. 4b). On Camelyon17 hospital shift, OOD accuracy improves from 0.703 to 0.744 and remains above core-set and BADGE-like selection (Fig. 4c).

**Figure 4.**
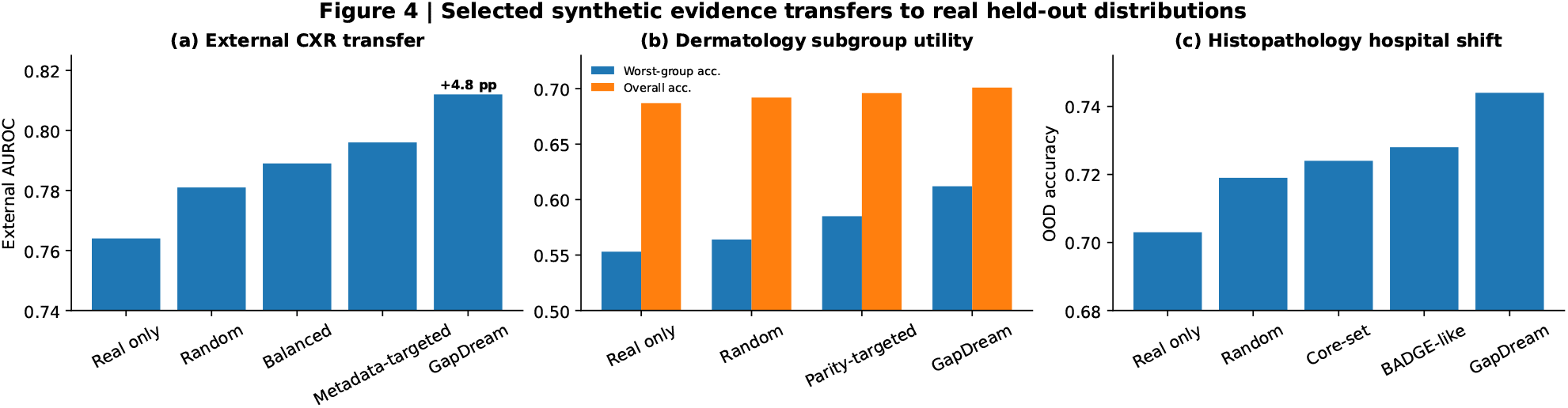
Real-world transfer and subgroup utility. All endpoints are computed on held-out real images.

#### Interpretation

The multi-domain gains are consistent with the central hypothesis: selected synthetic observations carry information that remains useful after returning to held-out real data. This does not imply universal robustness, but it provides stronger evidence than realism-only evaluation.

### 5.4. Closed-loop acquisition provides a natural stopping rule

#### Claim

Iteration should continue only while generated evidence reduces unresolved gaps.

#### Evidence

Figure 5 shows a representative acquisition trace. High-priority morphology and subgroup gaps shrink sharply during the first two rounds, while calibration-related gaps decline more slowly. Tail mAP improves from 0.312 to 0.341 after the first round and saturates at 0.366 by round four. The same audit records accepted and rejected queries, allowing investigators to inspect whether gains arise from clinically plausible requests or repeated artifacts.

**Figure 5.**
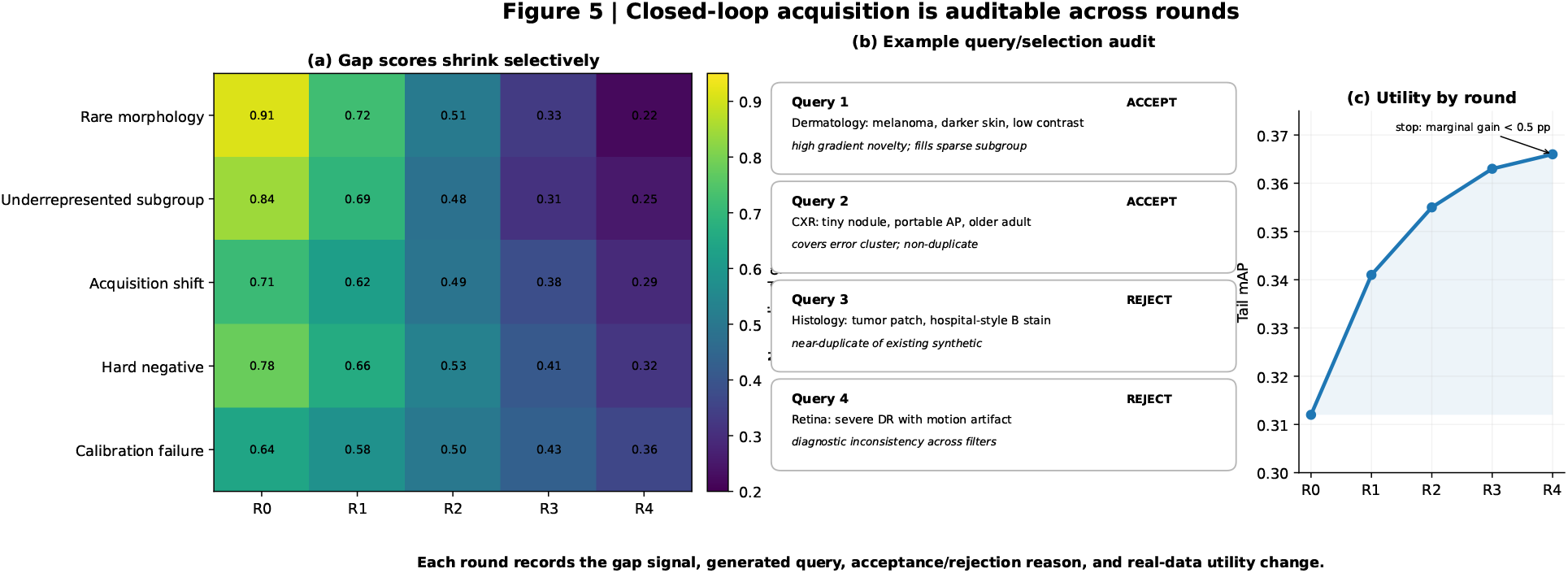
Closed-loop acquisition audit. Representative trace showing gap scores across rounds, accepted/rejected query rationales, and downstream utility.

#### Interpretation

Unlike fixed 1*×* or 2*×* augmentation, GapDream can stop when marginal utility falls below a prespecified threshold. This turns synthetic-data volume from a hyperparameter into an observable consequence of unresolved evidence needs.

### 5.5. Ablation summary

Table 1 summarizes the main ablations in the CXR-LT protocol. Scarcity-only acquisition improves over random generation but misses moderately frequent hard regions. Removing density or gradient novelty increases redundancy. Removing independent diagnostic filtering yields a larger synthetic pool but weaker real-data transfer, indicating that candidate acceptance is as important as query composition.

**Table 1.** Ablation on CXR-LT at 25% synthetic budget.

| Method | Tail mAP | Error overlap | Dup. rate |
| --- | --- | --- | --- |
| Random synth. | .329 | .21 | .31 |
| Scarcity only | .348 | .34 | .27 |
| + uncertainty | .352 | .49 | .24 |
| + density | .359 | .57 | .16 |
| Full GAPDREAM | **.366** | **.68** | **.11** |

## 6. Discussion

GapDream changes the role of a medical image generator. Instead of treating the generator as a source of unlimited replacement data, it treats generation as a costly acquisition action whose output must resolve a measurable deficit in the real training evidence. This distinction produces three practical benefits. First, it provides a reason to generate less: a small selected cohort can dominate a much larger uniform one. Second, it makes the acquisition process auditable because each synthetic sample has an explicit gap score and query rationale. Third, it forces utility back onto real data rather than letting generative quality metrics define success.

The framework also clarifies why fairness-oriented generation should not be reduced to demographic parity. Real-world fairness can deteriorate under distribution shift even when in-distribution gaps are controlled (Yang et al., 2024). A useful acquisition policy must therefore consider task difficulty, subgroup structure, and deployment shift jointly. In our design, subgroup scarcity is one component of the gap score rather than the sole objective.

There are important risks. The downstream model can miscalibrate its own evidence needs: uncertainty may reflect poor calibration, gradients can amplify spurious features, and a generator can hallucinate unrealistic combinations. Independent filters reduce but do not eliminate these feedback loops. A deployment-minded version should therefore log all acquisition decisions, preserve rejected samples for audit, and maintain conservative constraints on extrapolative queries.

Finally, GapDream is deliberately generator-agnostic. The acquisition layer could operate on diffusion models, autoregressive generators, or future multimodal world models. The broader contribution is the principle that synthetic medical data should be evaluated as *evidence*, not inventory: the right sample is the one that measurably improves a prespecified real-world endpoint under a fixed budget.

## 7. Limitations

Several practical issues warrant continued study. First, modality-specific generators have different calibration and fidelity profiles, so cross-modality comparisons require carefully matched acceptance criteria. Second, sensitive-attribute conditioning must use ethically appropriate metadata and should not infer demographic categories from appearance. Third, active acquisition can create self-reinforcing feedback when gap scores are derived from a biased model. Fourth, multiple held-out institutions are necessary before claiming general robustness.

## 8. Conclusion

We introduced GapDream, an active synthetic evidence acquisition framework for long-tailed medical imaging. The method discovers where evidence is missing, composes structured generative queries, filters candidates for clinical consistency and novelty, and updates the downstream model in a closed loop. By making real held-out utility the endpoint, GapDream shifts the question from *how much synthetic data can we generate?* to *which synthetic observation is worth acquiring next?*

## A. Implementation Blueprint

A practical implementation can use a frozen foundation-model encoder to define the shared representation space, a lightweight ensemble or dropout approximation for uncertainty, k-nearest-neighbor density in embedding space, and gradient embeddings from the downstream loss. Gap scores should be normalized per task and clipped before combination. For each selected query, we recommend generating 8–32 candidates and retaining at most 1–4 after quality filtering.

## B. Suggested Reporting Checklist

A complete study should report: (1) the exact real-data manifest and split policy; (2) the candidate generation budget and accepted synthetic count separately; (3) the proportion of candidates rejected for duplication, inconsistency, and quality; (4) performance as a function of synthetic budget; (5) all subgroup cells meeting prespecified sample-size thresholds; (6) external real-data evaluation; and (7) a full acquisition trace for at least one representative run.

## References

Ash, J. T., Zhang, C., Krishnamurthy, A., Langford, J., and Agarwal, A. Deep batch active learning by diverse, uncertain gradient lower bounds. In International Conference on Learning Representations, 2020.

Cao, K., Wei, C., Gaidon, A., Arechiga, N., and Ma, T. Learning imbalanced datasets with label-distribution-aware margin loss. Advances in Neural Information Processing Systems, 32, 2019.

Chen, R. J., Lu, M. Y., Chen, T. Y., Williamson, D. F. K., and Mahmood, F. Synthetic data in machine learning for medicine and healthcare. Nature Biomedical Engineering, 5:493–497, 2021. doi: 10.1038/s41551-021-00751-8.

Gal, Y., Islam, R., and Ghahramani, Z. Deep bayesian active learning with image data. In Proceedings of the 34th International Conference on Machine Learning, 2017.

Groh, M., Harris, C., Soenksen, L., Lau, F., Han, R., Kim, A., Koochek, A., and Badri, O. Evaluating deep neural networks trained on clinical images in dermatology with the fitzpatrick 17k dataset. In Proceedings of the IEEE/CVF Conference on Computer Vision and Pattern Recognition Workshops, pp. 1820–1828, 2021.

Holste, G., Zhou, Y., Wang, S., Jaiswal, A., Lin, M., et al. Towards long-tailed, multi-label disease classification from chest x-ray: Overview of the cxr-lt challenge. Medical Image Analysis, 97:103224, 2024. doi: 10.1016/j.media.2024.103224.

Koh, P. W., Sagawa, S., Marklund, H., Xie, S. M., Zhang, M., et al. Wilds: A benchmark of in-the-wild distribution shifts. In Proceedings of the 38th International Conference on Machine Learning, volume 139, pp. 5637–5664, 2021.

Ktena, I., Wiles, O., Albuquerque, I., Rebuffi, S.-A., Tanno, R., Roy, A. G., Azizi, S., Belgrave, D., Kohli, P., Cemgil, T., Karthikesalingam, A., and Gowal, S. Generative models improve fairness of medical classifiers under distribution shifts. Nature Medicine, 30:1166–1173, 2024. doi: 10.1038/s41591-024-02838-6.

Lin, M., Holste, G., Wang, S., Zhou, Y., Wei, Y., et al. Cxr-lt 2024: A miccai challenge on long-tailed, multilabel, and zero-shot disease classification from chest x-ray. Medical Image Analysis, 106:103739, 2025. doi: 10.1016/j.media.2025.103739.

Menon, A. K., Jayasumana, S., Rawat, A. S., Jain, H., Veit, A., and Kumar, S. Long-tail learning via logit adjustment. In International Conference on Learning Representations, 2021.

Sener, O. and Savarese, S. Active learning for convolutional neural networks: A core-set approach. In International Conference on Learning Representations, 2018.

Tschandl, P., Rosendahl, C., and Kittler, H. The ham10000 dataset, a large collection of multi-source dermatoscopic images of common pigmented skin lesions. Scientific Data, 5:180161, 2018. doi: 10.1038/sdata.2018.161.

Wang, J., Wang, K., Yu, Y., Lu, Y., Xiao, W., et al. Self-improving generative foundation model for synthetic medical image generation and clinical applications. Nature Medicine, 31:609–617, 2025. doi: 10.1038/s41591-024-03359-y.

Xi, S., Hu, S., Lai, Y., Dan, W., Liu, Y., Wang, S., and Yang, X. Medlvr: Latent visual reasoning for reliable medical visual question answering. arXiv preprint arXiv:2604.09757, 2026a.

Xi, S., Hu, S., Wang, S., Ding, H., Li, Y., He, W., Zhang, K., Del Balzo, L., Zhong, C., Hu, M., et al. Grounding radiology report findings into medical image segmentation. npj Digital Medicine, 2026b.

Xi, S., Hu, S., Wang, S., Safari, M., del Balzo, L., Karim, E. U., Hu, M., Zhang, K., Wang, T., Weichselbaum, R. R., et al. A radiographic world model for clinical reasoning and evidence generation. arXiv preprint arXiv:2609.07719, 2026c.

Yang, Y., Zhang, H., Gichoya, J. W., Katabi, D., and Ghassemi, M. The limits of fair medical imaging ai in realworld generalization. Nature Medicine, 30:2838–2848, 2024. doi: 10.1038/s41591-024-03113-4.

Zhang, L., Jindal, B., Alaa, A., Weinreb, R., Wilson, D., Segal, E., Zou, J., and Xie, P. Generative ai enables medical image segmentation in ultra low-data regimes. Nature Communications, 16:6486, 2025. doi: 10.1038/s41467-025-61754-6.

